# FUS controls the processing of snoRNAs into smaller RNA fragments that can regulate gene expression

**DOI:** 10.1101/409250

**Authors:** Patrycja Plewka, Michal Wojciech Szczesniak, Agata Stepien, Marek Zywicki, Andrzej Pacak, Martino Colombo, Izabela Makalowska, Marc-David Ruepp, Katarzyna Dorota Raczynska

**Author notes:** To whom correspondence should be addressed: Katarzyna Dorota Raczynska, Department of Gene Expression, Institute of Molecular Biology and Biotechnology, Adam Mickiewicz University, Wieniawskiego 1, 61-712 Poznan, Poland.

## Abstract

FUS is a multifunctional protein involved in many steps of RNA metabolism, including transcription, splicing, miRNA processing and replication-dependent histone gene expression. In this paper, we show for the first time that FUS binds and negatively regulates the levels of a subset of snoRNAs in cells. Scanning of available human small RNA databases revealed the existence of smaller RNA fragments that can be processed from FUS-dependent snoRNAs. Therefore, we suggest that FUS mediates the biogenesis of snoRNA-derived small RNAs, called sdRNAs. Further *in silico* approaches enabled us to predict putative targets of selected FUS-dependent sdRNAs. Our results indicate that sdRNAs may bind to different regions of target mRNAs as well as to noncoding transcripts and influence the posttranscriptional level or translation of these targets.

**SIGNIFICANCE STATEMENT:** RNA metabolism is orchestrated by a complex network of RNA-protein interactions and involves various classes of RNA molecules. Small nucleolar RNAs (snoRNAs) are commonly considered essential components of the ribosome biogenesis pathway. However, recent studies have revealed that snoRNAs can also be fragmented into small entities called snoRNA-derived RNAs (sdRNAs), which have been linked to multiple cancer types and thus may serve as next-generation prognostic or diagnostic biomarkers. In this paper, a multifunctional protein, FUS, was shown to be involved in the biogenesis of snoRNA-derived fragments. Furthermore, we combined bioinformatic analyses with complementary experimental approaches to elucidate the role of FUS-dependent sdRNAs in gene expression regulation. Our findings reveal the considerable regulatory potential of this new class of small noncoding RNAs.

## INTRODUCTION

Small nucleolar ribonucleoproteins (snoRNPs) are nucleolus-localized complexes consisting of small nucleolar RNAs (snoRNAs) associated with highly conserved core proteins. snoRNPs are grouped into two major classes, referred to as “box C/D” and “box H/ACA” snoRNPs, depending on the presence of “C/D” or “H/ACA” sequence motifs in the snoRNAs. Most snoRNPs function in precursor rRNA (pre-rRNA) processing by introducing sequence-specific modifications guided by snoRNAs. C/D snoRNPs are responsible for site-specific 2’-*O*-ribose methylation, while H/ACA snoRNPs catalyze the isomerization of uridine to pseudouridine. Moreover, members of another subset of snoRNPs - Cajal body-specific snoRNPs (scaRNPs) - modify the snRNA component of U snRNPs. Two enzymes are responsible for the catalytic activity of snoRNPs: the 2’-*O*-methyltransferase fibrillarin and the pseudouridine synthase dyskerin (DKC1) (1–3). Interestingly, a subset of snoRNAs that pair with rRNAs does not direct rRNA modification but rather acts as molecular chaperones required for correct folding of pre-rRNAs (reviewed in (4)). Furthermore, in higher eukaryotes, many “orphan” snoRNAs have been discovered. These snoRNAs contain either the C/D box or H/ACA box motif but do not contain an rRNA antisense region, suggesting that they may play different roles in the cell (1).

The snoRNP protein core includes four proteins. Box C/D snoRNPs consist of fibrillarin, NOP56, NOP58 and a 15.5 kDa protein; box H/ACA snoRNPs consist of dyskerin, GAR1, NHP2 and NOP10 (3, 5). The assembly of snoRNPs occurs entirely in the nucleus and is a complex multiple-step process that requires numerous transient assembly factors not found in mature, catalytically active snoRNP complexes. These assembly factors ensure the efficiency, specificity and quality control of snoRNP production. Assembly occurs in three major stages: i) formation of a protein-only complex that contains particular core proteins and assembly factors, ii) incorporation of nascent snoRNAs, and iii) release of the assembly factors and activation of snoRNP catalytic activity (6, 7). The formation of the complete core snoRNP complex is required for its nucleolar localization (8).

Most vertebrate snoRNAs are encoded within introns of pre-mRNAs, and the assembly of box C/D snoRNPs is generally dependent on splicing (9, 10). In contrast, the assembly of box H/ACA snoRNPs is believed to occur on nascent pre-mRNAs and is connected to RNA polymerase II (RNAP2) transcription. This connection is mediated by nuclear assembly factor 1 (NAF1), which interacts with the C-terminal domain of the large subunit of RNAP2 and may recruit H/ACA core proteins to newly synthesized snoRNAs (11). Nevertheless, for both classes of snoRNPs, splicing of the host pre-mRNA is essential for providing linear precursor snoRNA, which is generated from debranched intron lariats and further processed by exonucleases: XRN1/2 family members from the 5’ end, and RNA exosome with NEXT complex from the 3’ end (reviewed in (7)).

Accumulating evidence indicates that many snoRNAs are processed into shorter functional forms with lengths of 19-40 nucleotides (reviewed in (4, 12)). snoRNA-derived RNAs (sdRNAs) have been identified in animals (human (13–19)), rodents (15, 20), Drosophila, and chicken (15)) as well as in plants (*Arabidopsis thaliana*) (15), fission yeast (*Saccharomyces cerevisiae*) (15), protozoa (*Giardia lamblia*) (21) and the Epstein-Barr virus (22). The mechanism of sdRNA generation is, as of yet, not completely understood. Many H/ACA sdRNAs are produced in a Dicer-dependent manner and associate with Argonaute (AGO) proteins (13–16, 23, 24). In contrast, C/D box-derived small RNAs are not efficiently incorporated into AGO2 proteins, and most originate from the termini of mature snoRNAs and hence carry “C” and “D” box motifs (25, 26).

Functionally, many sdRNAs exhibit microRNA-like properties in posttranscriptional gene silencing activity (13–16). However, numerous noncanonical functions, such as mediating mRNA editing and splicing, have recently been ascribed to sdRNAs (12, 24, 27, 28). Interestingly, a Dicer-dependent sdRNA was identified in the cytoplasmic fraction of *Giardia lamblia* cells, and its function in translational regulation was experimentally validated (21). The presence of sdRNA (originating from both box C/D and box H/ACA snoRNAs) was also demonstrated in *S. cerevisiae*. Interestingly, sdRNA processing events were most prominent under nonoptimal conditions for yeast growth, including UV irradiation, oxygen deprivation, high/low pH exposure or culture in medium with no amino acids or sugars. This response to environmental conditions may, therefore, indicate that sdRNA processing plays a crucial role in the regulation of stress-dependent metabolism (29). Importantly, these sdRNAs have been copurified with yeast ribosomes, implicating a novel yet-undiscovered regulatory role of sdRNAs in ribosome biosynthesis (29). In addition, sdRNAs have been implicated in human cancer, and according to high-throughput analyses performed by the Chen group, sdRNAs can be prevalent molecular markers across multiple types of human cancer (30).

In this paper, we show for the first time that fused in sarcoma (FUS) protein is involved in the biogenesis of small RNAs derived from mature snoRNAs in human cells. FUS belongs to the FET family of proteins, which includes three highly conserved, abundant, ubiquitously expressed RNA-binding proteins: FUS, EWS and TAF15 (31). FUS is a nucleocytoplasmic shuttling protein predominantly present in the nucleus but is also found in the cytoplasm and was suggested to participate in mRNA transport and translation (32, 33). The protein also binds to DNA (single-stranded DNA (ssDNA) as well as double-stranded DNA (dsDNA)), facilitating DNA annealing and D-loop formation and thus mediating genomic maintenance, DNA recombination and DNA repair (34–36). FUS regulates several key steps in RNA metabolism, including transcription, splicing and alternative splicing (reviewed in (37, 38)). Additionally, FUS is involved in replication-dependent histone gene expression (39) and may play a role in miRNA processing as a component of the large Drosha microprocessor complex (40). Notably, several FUS mutations have been found in familial forms of amyotrophic lateral sclerosis (ALS) and frontotemporal lobar degeneration (FTLD), implicating a role for this protein in neurodegenerative diseases (31, 38, 41–44).

Here, we show that FUS can bind snoRNAs in human cells. Using RNA immunoprecipitation with anti-FUS antibodies followed by high-throughput RNA sequencing (RIP-seq), we identified all three classes of snoRNAs in the immunoprecipitated fraction. The interaction of FUS with snoRNA fragments was further confirmed by an electrophoretic mobility shift assay (EMSA) and a double filter binding assay. Surprisingly, we observed that FUS negatively influences the level of mature snoRNAs in cells, although the splicing efficiency of snoRNA-hosting introns was not altered. Interestingly, scanning of available human small RNA databases revealed the existence of sdRNAs with lengths of 19-33 nucleotides that can be derived from a subset of snoRNAs. Therefore, we propose that FUS does not affect snoRNA biogenesis but rather competes with snoRNP proteins to regulate the synthesis of FUS-dependent sdRNAs. Further *in silico* approaches enabled the prediction of putative targets of our identified sdRNAs. Our results show that FUS-dependent sdRNAs might regulate gene expression by controlling transcripts level or translation efficiency.

## RESULTS

### FUS directly interacts with snoRNAs in human cells

Total and nuclear extracts from HEK293T cells overexpressing FLAG-FUS and FLAG-EBFP (negative control) (Supplementary Fig. S1A) were subjected to immunoprecipitation with anti-FLAG antibodies followed by isolation of RNA from the immunoprecipitated fractions. Coprecipitated RNAs were analyzed by high-throughput sequencing. The analysis of small noncoding RNAs bound by FUS revealed a large fraction of snoRNAs belonging to three classes of snoRNAs: box C/D snoRNAs (snord), H/ACA snoRNAs (snora) and small Cajal body-specific RNAs (scaRNAs). FUS-bound snoRNAs were enriched in fractions immunoprecipitated from nuclear extracts, with a distribution resembling that of endogenous snoRNAs in cells—approximately 67% snord, 27% snora and 6% scaRNA (Fig. 1A, Supplementary Table S1). This result suggests that FUS binds to snoRNAs in a class-independent manner.

**Figure 1.**
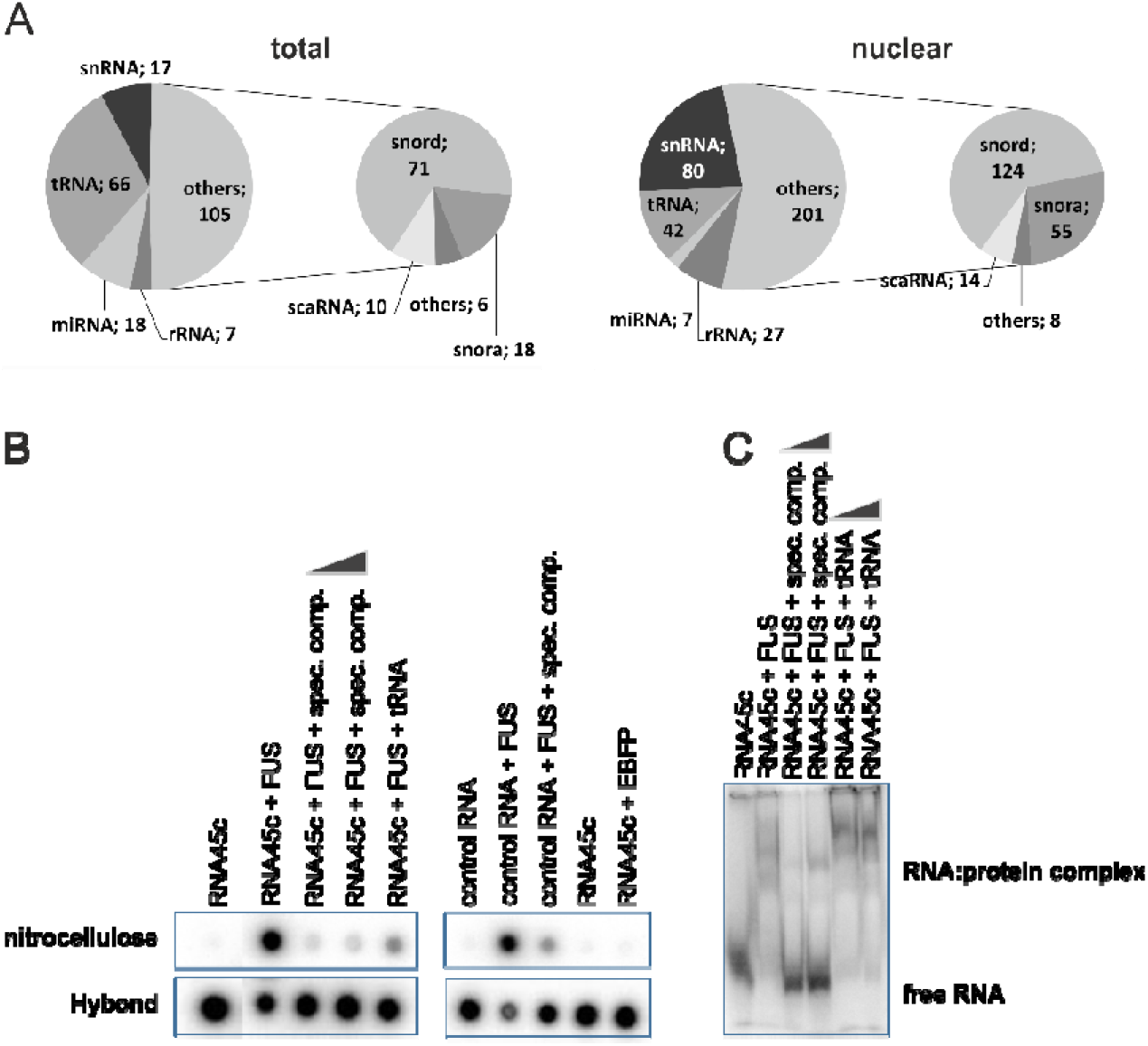
*A) Distribution of FUS-bound small noncoding RNAs identified in fractions immunoprecipitated from total and nuclear extracts. The numbers represent noncoding RNAs with ≥4-fold enrichment relative to their expression in the control sample. B) The double filter binding assay and (C) EMSA assay confirmed that FUS can directly interact with snoRNAs. RNA fragments encompassing the C/D motifs of snord45c (RNA45c) and the positive control sequence (control RNA) were subjected to interaction with purified recombinant FLAG-FUS and FLAG-EBFP (as the negative control). The binding specificity was tested by adding specific (unlabeled RNA fragments encompassing the C/D motifs of snord45c) and nonspecific (tRNA) competitors*.

To confirm that FUS can directly interact with snoRNAs, we performed a double filter binding assay (Fig. 1B) and an EMSA assay (Fig. 1C). Isotope-labeled RNA fragments encompassing the C/D motifs of snord45c and a control sequence encompassing the GGUG motif recognized by FUS (positive control, as described by (45)) were incubated with recombinant FLAG-FUS and FLAG-EBFP (negative control). The binding specificity was assessed by the addition of specific (unlabeled RNA) and nonspecific (tRNA) competitors. As shown in Fig. 1B and C, we confirmed that FUS can directly interact with snoRNAs.

### FUS negatively regulates the level of mature snoRNAs

As FUS interacts directly with snoRNAs, we addressed the question of whether FUS can affect the level of mature snoRNAs in cells. We isolated total RNA from control cells and cells with FUS overexpression (FUS OE), inducible FUS knockdown (FUS KD) and FUS knockout (FUS KO) (Supplementary Fig. S1). cDNA synthesized in a coupled polyadenylation reverse transcription reaction was used as the template for qPCR with a universal reverse primer and a snoRNA-specific forward primer (39). The level of several mature snoRNAs changed in a manner inversely related to the amount of FUS in the cell, suggesting that FUS acts as a negative regulator of mature snoRNAs (Fig. 2A).

**Figure 2.**
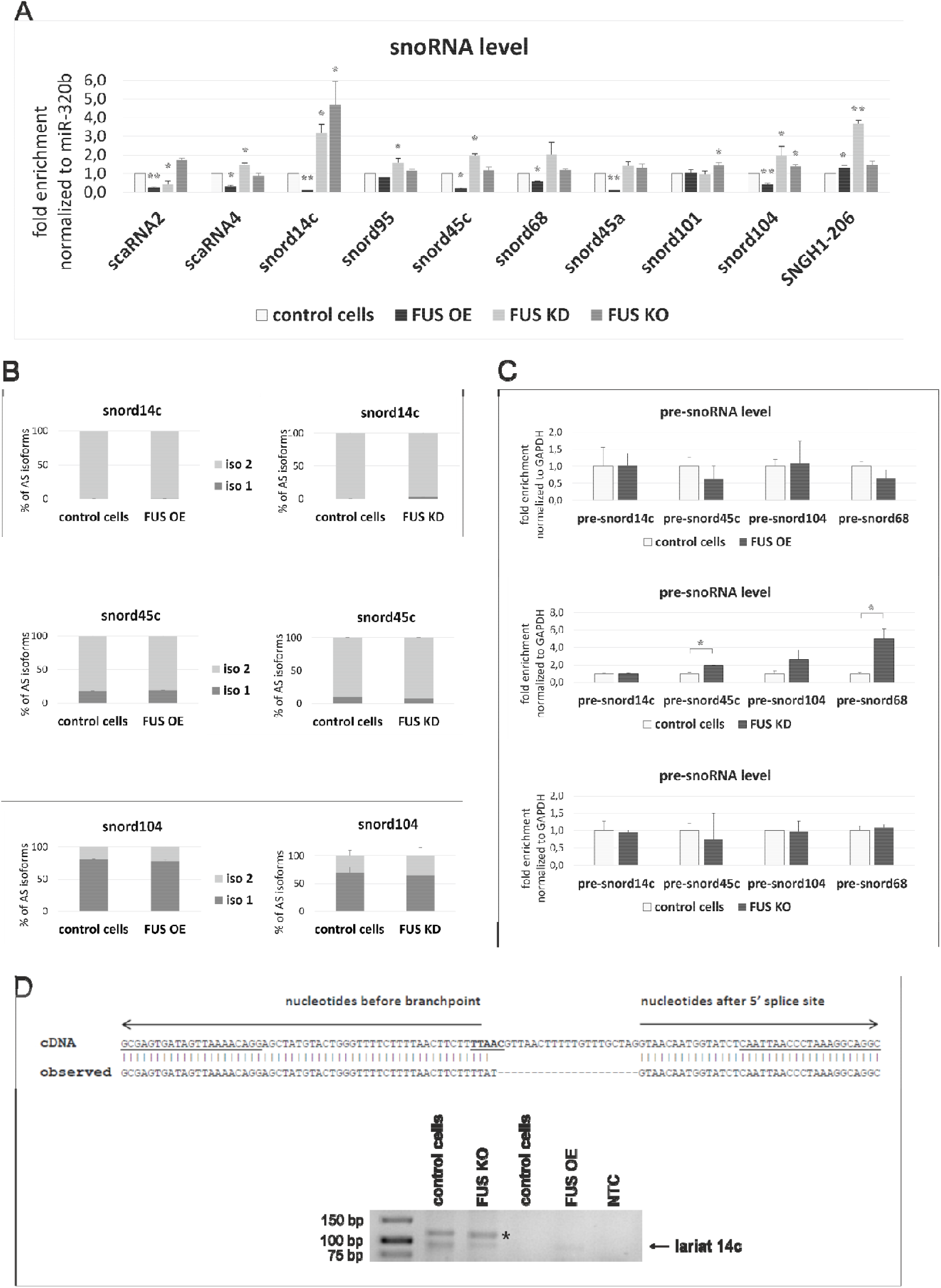
*FUS affects the level of mature snoRNAs. A) The effect of FUS overexpression (FUS OE), FUS knockdown (FUS KD) and FUS knockout (FUS KO) on the level of selected mature snoRNAs was analyzed by RT-qPCR. B) The effect of FUS overexpression (FUS OE) and FUS depletion (FUS KD) on the splicing of introns encoding snoRNA genes was analyzed by RT-qPCR. Primers were designed to amplify alternative isoforms (iso1 (spliced isoform) and iso2 (unspliced isoform)). C) The effect of FUS overexpression (FUS OE) and FUS depletion (FUS KD and FUS KO) on the level of snoRNA precursors was analyzed by RT-qPCR. The error bars indicate the standard deviations (SDs) of three biological replicates. P-values were calculated using Student’s t-test, and statistical significance is represented as follows: *p≤ 0.05 and **p ≤ 0.01. D) (Lower panel) RT-PCR using outward-facing primers was performed to amplify lariat 14c in FUS KO, FUS OE, control cells and nontemplate control (NTC). The star indicates an nonspecific product. (Upper panel) sequencing of the lariat 14c product revealed a 14-nucleotide gap between the branch point and the 5’ splice site. The sequence of the predicted branch point is in bold and underlined, and the sequences of primers are underlined*.

In vertebrates, the vast majority of box C/D and box H/ACA snoRNAs are encoded within intronic sequences of pre-mRNAs, and the maturation of these snoRNAs depends on splicing (9, 10). Therefore, we asked whether FUS affects snoRNA biogenesis by changing the splicing profiles of snoRNA-encoding introns. To test this hypothesis, we analyzed the ratio between spliced and unspliced isoforms of snoRNA-hosting introns. We could not find any changes in the splicing efficiency of these introns in cells with either FUS OE or FUS depletion (Fig. 2B). Furthermore, we tested the levels of snoRNA precursors in cells with FUS OE, cells with FUS KD and cells with FUS KO. Again, we could not observe significant changes, except for elevated levels of pre-snord45c and pre-snord68 in cells with FUS KD (Fig. 2C). This finding suggests that FUS predominantly influences snoRNA levels at later steps of snoRNA biogenesis.

Recent studies in human cells have identified numerous circular intronic long noncoding RNAs as well as long noncoding RNAs produced from introns with two imbedded snoRNA genes (46, 47). In addition, stable intronic sequence RNAs (sisRNAs) have been described in *Xenopus laevis* (48). These sisRNAs can be linear or can form debranching-resistant lariat structures and can accumulate in both the nucleus and the cytoplasm. Interestingly, cytoplasmic lariat sisRNAs originate from small introns (<200 nt) and can encode snoRNA genes (48). Considering these data, we decided to test whether snoRNAs can be trapped by circular lariat introns that are derived from spliced introns and escape debranching. Indeed, when we amplified cDNA by PCR using outward-facing primers, we detected a product with the expected size of the lariat intron encoding snord14c (lariat 14c). The lariat 14c product was further cloned, and its origin was confirmed by sequencing. Similarly, as described in (48), we determined that although the sequence of this product mapped to the region between the 5’ and the 3’ ends of the intron, one end mapped precisely to the 5’ splice site, whereas the other end mapped to a region 14 nucleotides upstream of the 3’ splice site, at the presumed branchpoint (Fig. 2D, upper panel). This result suggests that lariat 14c was derived from splicing, escaped debranching and was stabilized in the form of a lariat without a tail. Interestingly, the expression level of lariat 14c was downregulated in FUS KO cells and upregulated in FUS OE cells; thus, the formation of lariat 14c could be induced by FUS (Fig. 2D, lower panel). We assume that lariat formation might serve to block mature snoRNA production or protect snoRNA from further processing into sdRNA (see Discussion).

### FUS induces the processing of mature snoRNAs into smaller RNA fragments—sdRNAs

Within the last few years, many reports have been published concerning sdRNAs and their origin from snoRNAs (reviewed in (4, 12)). sdRNA biogenesis should lead to a decrease in mature snoRNAs in the cell. Therefore, we addressed the question of whether FUS could be involved in the generation of small RNA fragments from selected mature snoRNAs. We examined available high-throughput sequencing data for small RNAs isolated from three different human cell lines (HEK293T cells (SRR1586016), neuroblastoma cells (SRR3931966) and whole blood (SRR2728234)). We tried to identify sdRNAs that could be derived from several FUS-dependent snoRNAs—snord104, snord45c, snordD14c and snord68 (Fig. 2A)—and found that small RNAs could be mapped to these snoRNAs in all three libraries. As shown in Fig. 3A, with the exception of snord68, small RNAs mapped typically to the sequences in two clusters: one near the 3’ end of the snoRNA, with reads having a quite homogeneous 3’ end; and the other close to the 5’ end with a homogeneous 5’ end (25). Our *in silico* approach enabled us to identify putative short sdRNAs (with lengths of <25 nt) embedded in snord45c, snord14c and snord104. In addition to short, miRNA-like reads, many longer reads mapped to the snoRNAs (with lengths of ≥25 nt) embedded in snord68 and snord104 (Fig. 3A).

**Figure 3.**
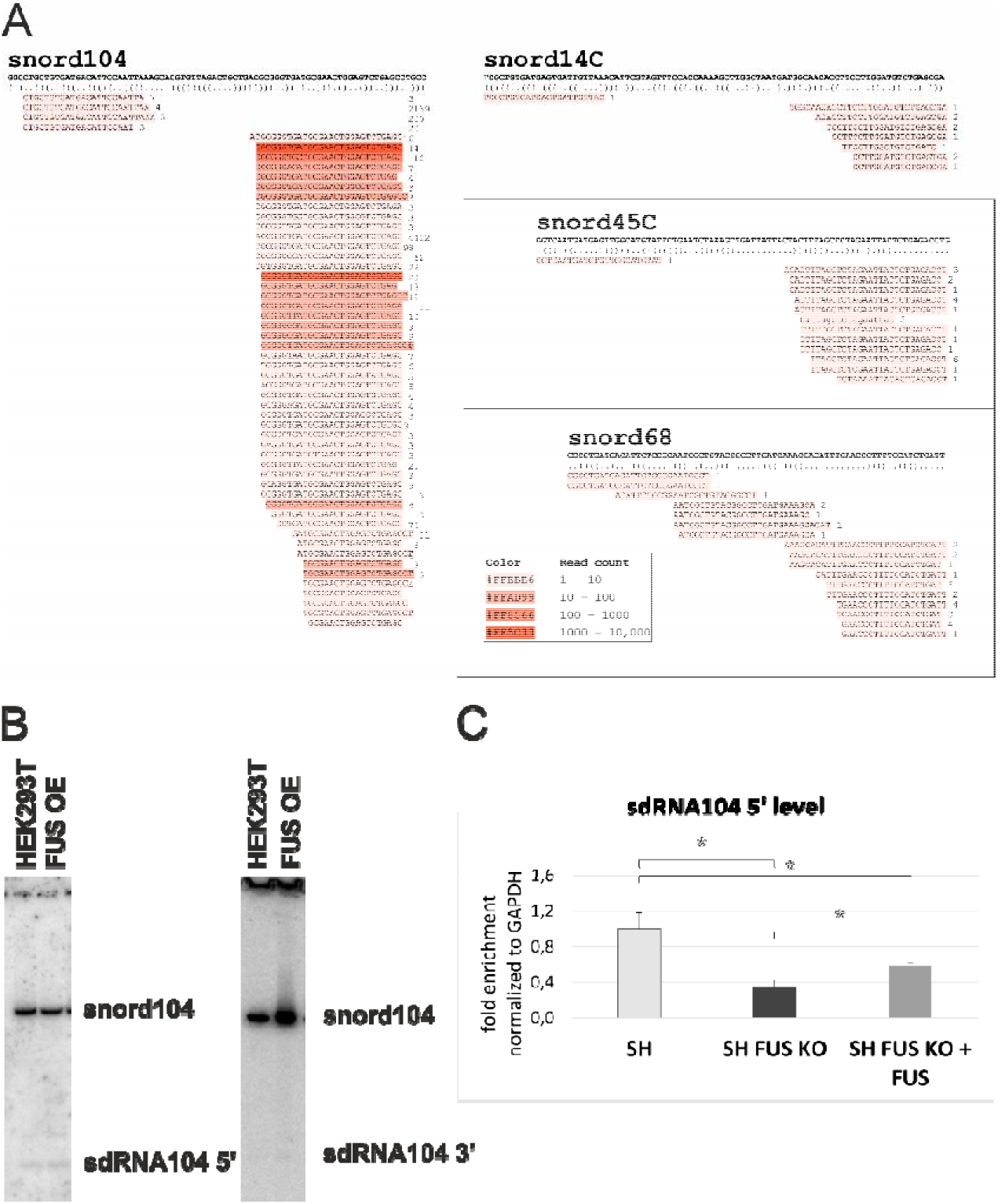
*snoRNAs can be processed into sdRNAs, and FUS controls this process. A) Scanning of human small RNA databases (derived from cultured HEK293T cells, human neuroblastoma cells and whole blood) revealed the existence of sdRNAs with lengths of 19-33 nt, which can be derived from select snoRNAs. The numbers on the right indicate the numbers of reads. Secondary structures were predicted with RNAfold (http://rna.tbi.univie.ac.at/cgi-bin/RNAfold.cgi): minimum free energy prediction and optimal secondary structure. B) Northern blot hybridization confirmed the existence of sdRNA104s derived from the 5’ and 3’ ends of snord104 in cells. C) The level of sdRNA104 derived from the 5’ end of snord104 was analyzed by SL RT-qPCR in SH-SY5Y wild type cells (SH) and cells with FUS knockout (SH FUS KO) complemented by exogenous FUS (SH FUS KO + FUS). The error bars represent the SDs of three biological replicates. P-values were calculated using Student’s t-test, and statistical significance is represented as follows: *P ≤ 0.05*.

Next, we confirmed by two different methods the presence of sdRNA104 5’ (a small RNA derived from snord104 and processed from the 5’ end of snord104) in two different cell lines and showed that the level of this sdRNA depends on FUS expression. Firstly, we performed Northern blot hybridization using a probe targeting sdRNA104 embedded in either the 5’ end or 3’ end of snord104. Interestingly, we detected a similar signal for both sdRNA104 probes (Fig. 3B), although the number of reads from high-throughput sequencing was higher for sdRNA104 derived from the 3’ end of snord104 than for sdRNA104 derived from the 5’ end of snord104 (Fig. 3A). Secondly, we performed stem-loop reverse transcription PCR (SL RT-PCR), which showed that the expression level of sdRNA104 5’ was significantly decreased in neuroblastoma SH-SY5Y cells with FUS KO and was partially restored in FUS KO cells complemented with exogenous FUS (Fig. 3C). These results suggest that FUS induces sdRNA104 processing from the 5’ end of snord104. No effect on sdRNA104 expression was observed in SH-SY5Y cells with transient overexpression of FUS (not shown). However, notably, we did not observe a significant increase in FUS protein levels in these cells (Supplementary Fig. S2), probably due to FUS autoregulation (49). Unfortunately, we could not detect any other FUS-dependent sdRNAs by Northern blot hybridization, most likely due to the very low sdRNA levels in cells. Additionally, the specificity of SL RT-PCR enabled us to reliably detect and quantify only sdRNAs originating from the 5’ termini of mature snoRNAs.

### sdRNAs can regulate the level of transcripts and translation of mRNAs

The function of many sdRNAs has not yet been elucidated. Some have been shown to act like miRNAs, as they suppress target gene expression through the inhibition of translation or the acceleration of mRNA degradation after complementary Watson-Crick base pairing with their target transcripts (50). Other sdRNAs were reported to bind to introns/exons thereby influencing the splicing profile of the targeted pre-mRNAs (24, 27, 28). To further determine the function of FUS-dependent sdRNAs, we first sought their putative targets using Miranda software with either the whole transcriptome or only 3’ UTRs as the input. The criteria for sdRNA:RNA hybrid formation resemble those for miRNA:target hybridization. To support *in silico* target predictions, available datasets from the crosslinking immunoprecipitation (CLIP)-Seq experiment for AGO proteins were used.

Our *in silico* approach revealed that selected FUS-dependent sdRNAs have hybridization potential to the 3’ UTR of mRNAs. This binding, in turn, may impact translation efficiency. To evaluate the potential influence of sdRNAs on the expression of targeted transcripts, we assessed the effect of the interaction between sdRNA104 5’ and brain and reproductive organ-expressed protein (BRE) mRNA (ENSG00000158019) (Fig. 4A). We cloned the 3’ UTR region of the BRE transcript downstream of the luciferase (LUC) coding sequence. Then, the effect on luciferase synthesis was determined in FUS KO cells, in which we expect a lower level of sdRNA. Indeed, as shown in Fig. 4B, 48 h after plasmid transfection, we observed decreased luciferase activity. This result was confirmed by Western blot analysis using anti-BRE antibodies (Fig. 4C). We concluded that FUS-dependent sdRNAs might stimulate the translation or stabilize the binding of the BRE transcript to ribosomes.

**Figure 4.**
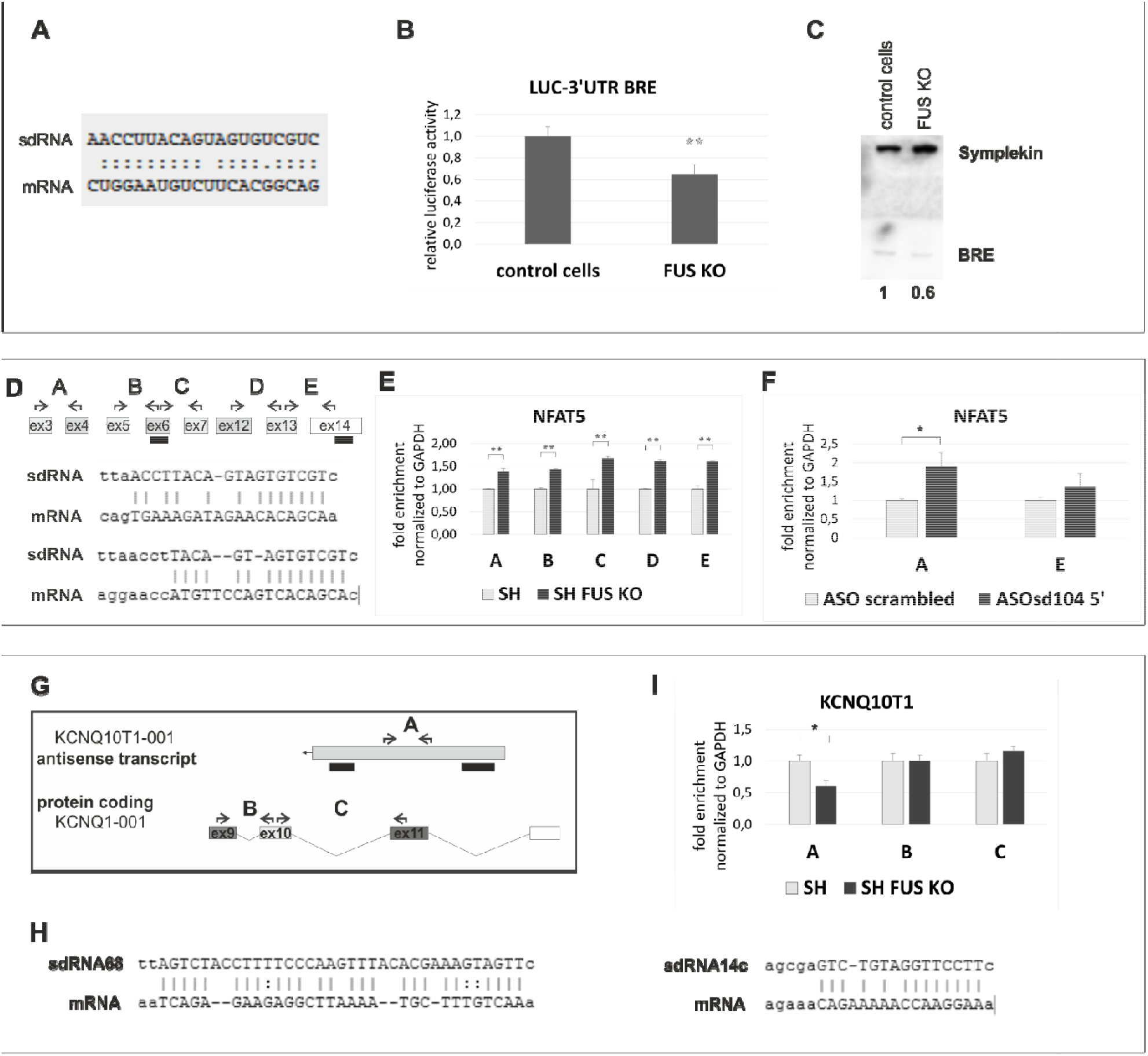
*sdRNAs are involved in the regulation of gene expression. A) sdRNA104 5’:BRE mRNA duplex. (B) Luciferase assays confirmed the positive effect of sdRNA on mRNA translation. The 3’UTR of BRE mRNA targeted by FUS-dependent sdRNA104 5’ was cloned downstream of the luciferase coding sequence, and its effect on translation was assessed in FUS KO cells. C) Western blot followed by immunodetection with anti-BRE and anti-Symplekin (loading control) antibodies was performed using protein extracts from control cells and FUS KO cells. The relative amount of BRE protein is listed below. D) (Upper panel) sdRNA104 5’ (indicated by black bars) can target NFAT5 mRNA. The primer pairs (A-E) were located as shown in the diagram. (Lower panel) sdRNA104 5’:NFAT mRNA duplexes. E, F) RT-qPCR was performed using a set of primer pairs designed as shown in the diagram. G) (upper panel), sdRNAs (indicated by the black bars) can target KCNQ10T1 antisense transcripts. H) The sdRNA:target duplexes are formed within a unique region that distinguishes noncoding RNA from mRNA transcribed from the same genomic region. I) RT-qPCR analysis was performed using a set of primers designed as shown in the diagram in (G). The error bars represent the SDs of three biological replicates. P-values were calculated using Student’s t-test, and statistical significance is represented as follows: *P ≤ 0.05 and **P ≤ 0.01*.

According to another bioinformatic prediction, FUS-dependent sdRNA104 derived from the 5’ end of snord104 can hybridize to different exons of NFAT5 mRNA (Fig. 4D). As FUS is involved in splicing and alternative splicing (37, 38), we first asked whether intron excision is altered in cells with FUS depletion. We designed primer pairs that hybridize to exons targeted by sdRNA104 5’ or to neighboring exons (Fig. 4D). Interestingly, the RT-qPCR results revealed that the level of total mRNA but not the splicing profile was altered. We observed similar changes in the transcript level regardless of the location of the primer pairs within the gene. NFAT5 mRNA level was upregulated in FUS KO cells (Fig. 4E). Moreover, the same effect was observed in cells in which we depleted sdRNA104 derived from the 5’ end of snord104 using an antisense oligonucleotide (ASO) strategy (Supplementary Fig. S3), suggesting a direct and negative influence of sdRNA104 on NFAT5 gene expression (Fig. 4F).

As shown in Fig. 4G, our *in silico* approach indicated that FUS-dependent sdRNA68, originating from snord68, might target KCNQ10T1-001 antisense transcripts encoded by the opposite strand of the KCNQ1 protein-coding gene. Interestingly, the same antisense KCNQ10T1 transcript might be targeted by sdRNA14c (Fig. 4G, H). The sdRNA:target duplexes are formed within the unique region that distinguishes noncoding RNA from mRNA transcribed from the same genomic region. The KCNQ10T1 antisense transcript (ENSG00000269821) encompasses intron 10, exon 11 and intron 11 of the KCNQ1 mRNA (ENSG00000269821), and the sdRNA-targeted regions within the antisense transcript correspond to intron 10 and intron 11 within the KCNQ1 mRNA (Fig. 4G). RT-qPCR analysis using a set of primers that specifically amplify the noncoding or coding transcript revealed that only antisense KCNQ10T1 transcript was significantly affected in FUS KO cells (Fig. 4I). The amount of KCNQ1 mRNA did not change under these conditions. One possible scenario assumes that the increased level of sdRNA14c, which is liberated from the lariat structure in FUS KO cells (Fig. 2D), induces the degradation of the antisense transcript. Alternatively, both of the short RNAs that can target the antisense KCNQ10T1 transcript—sdRNA68 and sdRNA14c—may exhibit activity, and the final level of the KCNQ10T1 transcript is a result of interplay between these two sdRNAs. However, the role of sdRNA68 in this regulation could not be described, as our experimental approaches could not determine sdRNA68 in cells.

## DISCUSSION

As suggested by Bratkovic and Rogelj, at least three reasons suggest that sdRNAs are not simply degradation products of snoRNAs: i) common processing patterns have been observed, ii) the processing extent of different snoRNAs may differ in the same cell line, and iii) posttranscriptional gene regulation activity has been confirmed for some sdRNAs (4). Our data show for the first time that in human cells, FUS mediates the biogenesis of a subset of sdRNAs, which are involved in the regulation of gene expression at different levels.

The mechanism of sdRNA production is still not fully elucidated. According to our results, the trimming of selected snoRNAs into sdRNAs is mediated by FUS, and this process involves mature snoRNAs. Mature snoRNAs are processed from removed and debranched introns by exonucleolytic activity, and we found that FUS did not affect the splicing profile of host genes (Fig. 2B). We suggest that instead, FUS predominantly affects the level of snoRNAs at later steps in snoRNP biogenesis.

The assembly of box C/D and box H/ACA snoRNPs operates via a multistep mechanism that requires both core proteins and assembly factors (6, 7). During the biogenesis of box C/D snoRNPs, a protein-only complex containing SNU13 and NOP58, in association with assembly factors such as the AAAC ATPase RUVBL1/2, NUFIP, ZNHIT3 and ZNHIT6, is preformed with assistance from the HSP90/R2TP complex and loaded on snoRNA in a cosplicing-dependent manner. Pre-snoRNP particles are then transported to Cajal bodies (CBs), where final processing occurs, and catalytically active snoRNPs are then transported to nucleoli. However, whether assembly factors leave pre-snoRNPs before or after their arrival in CBs is not clear. During the assembly of box H/ACA snoRNPs, two conserved proteins—NAF1 and SHQ1—are required for the stability of box H/ACA snoRNAs but are not part of the mature particles. In the final step of biogenesis, NAF1 is replaced by GAR1, which leads to the production of mature and functional H/ACA snoRNPs (6, 7). In mammalian cells, the SMN complex has been suggested to play a role in this exchange (51). We assume that FUS can interact with any of these proteins (as has been shown for SMN (52)) to impede the completion of snoRNP assembly. Alternatively, by such interactions, FUS can recruit or facilitate the action of other proteins involved in the exo- or endonucleolytic cleavage of snoRNAs into smaller fragments. As a third possibility, FUS could induce sdRNA production at the stage of snoRNP disassembly. However, regardless of whether FUS impedes snoRNP assembly or induces snoRNP disassembly, sdRNA production leads to decreased levels of mature snoRNAs/snoRNPs in cells.

An alternative hypothesis states that snoRNAs can be trapped by circular lariat introns derived from spliced introns, which escape debranching and are protected from further processing. The formation of such lariat introns containing snoRNAs could be mediated by FUS. Additionally, if so, an interaction between FUS and snoRNP proteins might be necessary. The possibility that this step is required to generate sdRNAs cannot be excluded. Alternatively, FUS could induce the generation of lariat structures to trap and block the maturation or processing of snoRNAs. Similar to the pathways described in *Arabidopsis thaliana*, where intron lariat RNAs inhibit miRNA biogenesis, this process could be another mechanism regulating snoRNA/sdRNA levels (53). The function of intronic RNAs is largely unknown; they may play a role in local gene transcription, sequester proteins or even act as repositories for snoRNAs in cells.

Our RIP-seq experiment identified a small fraction of tRNA-derived fragments (tRFs), so the possibility that FUS induces the synthesis of other small regulatory RNAs cannot be excluded. The biogenesis of most tRFs is incompletely described. The role of Dicer, DGCR8 or Drosha is suggested to be an exception, rather than a rule; however, enzymes involved in the liberation of tRFs from the 5’ and 3’ ends of mature tRNAs are still not fully known (54, 55). Thus, the generation of tRFs and sdRNAs might proceed similarly and might involve the FUS protein. Deeper understanding of the molecular mechanism underlying sdRNA biogenesis might, therefore, shed light on the processing of tRFs and other small RNAs mediated by FUS.

Interestingly, among our identified FUS-dependent sdRNA targets were transcripts highly expressed in the brain (The Human Protein Atlas; http://www.proteinatlas.org). For example, as a component of the BRCA1-A complex, BRE functions in DNA damage repair as well as in apoptosis prevention. BRE is thought to play a role in homeostasis or cellular differentiation in cells of neural, epithelial and germline origins. Moreover, recent research has further illuminated the importance of snoRNA fragments in cancer tissues. The Bangma and Jenster group observed that sdRNAs display strong differential expression and are massively upregulated in prostate cancer (56). Recently, high-throughput analyses performed by the Chen group allowed the formation of a comprehensive map of the sdRNA transcriptome across multiple human cancer types (30). Importantly, BRE and KCNQ10T1 are disease-related genes. Marked overexpression of BRE was detected in many tumors, suggesting that this gene promotes cell proliferation and local tumor growth (57, 58). The epigenetic status of KCNQ10T1 was shown to be correlated with colorectal carcinogenesis (59). In summary, sdRNAs and human diseases are linked. Moreover, FUS is implicated in neurodegeneration, tumors and cellular stress responses through errors in multiple steps of RNA processing. Therefore, our preliminary results showing a connection between FUS-dependent sdRNAs and disease-related genes are promising and might further indicate the role of FUS and FUS-dependent sdRNAs in oncogenic networks of select tumors.

## MATERIALS AND METHODS

Antibodies and general methods for cell culture and transfection, RNA isolation, cDNA preparation, PCR, protein isolation, immunoprecipitation, Western blot analysis, RNA library preparation and bioinformatic analysis are described in the Supplementary Materials and Methods. The original protocols are described below.

### Northern blot analysis

Northern blot analysis was performed as previously described (60). Briefly, total RNA (40 μg) was separated on 15% denaturing polyacrylamide gels containing 7 M urea. Electrophoresis was performed in 20 mM MOPS-NaOH buffer (pH 7.0), and RNA DecadeMarker (Thermo Fisher Scientific) was loaded as the lenght marker. After electrophoresis, RNA was transferred onto an Amersham Hybond-NX nylon membrane with a Trans-Blot SD Semi-Dry Transfer Cell (Bio-Rad) by applying 20 V for 1 h and then fixed using the EDC-mediated (1-ethyl-3-(3-dimethylaminopropyl) carbodiimide) chemical cross-linking method (2 h at 55°C). For the detection of snoRNAs and sdRNAs, 1 h prehybridization and 16 h hybridization steps were performed at 37°C in hybridization buffer (3.5% SDS, 0.375 M sodium phosphate dibasic, and 0.125 M sodium phosphate monobasic) containing DNA probes (Sigma-Aldrich) labeled at the 5’ end with [γ-^32^P]-ATP. Membranes were then washed twice in 2x SSC buffer containing 0.1% SDS to remove any unbound probe, and blots were exposed for three days to a phosphorimaging screen (Fujifilm) and scanned with a Fujifilm FLA5100 reader (Fujifilm).

### Electrophoretic mobility shift assay and double filter binding assay

RNA fragments encompassing the C/D motifs of snord45c and a control sequence (control RNA) encompassing the GGUG motif recognized by FUS (as described by (45)) were labeled by *in vitro* transcription in the presence of [⍰-^32^P]-UTP. Recombinant FLAG-FUS and FLAG-EBFP (as a negative control) proteins were overexpressed and purified from HEK293T cells. For the EMSA and double filter binding assay, 150 ng of protein was incubated with labeled RNA fragments (10 000 cpm) for 30 min at room temperature in buffer containing 20 mM Tris-HCl, pH 7.4; 50 mM KCl; and 10% glycerol. To test the binding specificity, unlabeled RNA fragments or tRNA (30 ng and 60 ng) were added to the samples 5 min before labeled RNA was added. All RNAs were denatured for 1 min at 95°C and cooled to RT before being added to the reaction. Samples were loaded on a 5% polyacrylamide gel, and electrophoresis was performed at 200 V and 15-20 mA in 0.5x TBE for approximately 2.5 h at 4°C. For the double filter binding assay, after incubation, samples were passed through Nitrocellulose/Amersham Hybond membranes using a vacuum. The gel and membranes were dried and exposed to a phosphorimaging screen (Fujifilm).

### Luciferase reporter assay

The 3’ UTR region of BRE mRNA was amplified by PCR with primers introducing *NheI* and *SalI* restriction sites and inserted into the pmirGLO vector after linearization by *NheI* and *SalI*, leading to the formation of the LUC-3’UTR BRE luciferase reporter plasmid. The PCR primer sequences used are available on request. Control cells and FUS KO cells (approximately 2x10^4^ cells/well) were cotransfected with 0.6 μg of the LUC-3’UTR BRE luciferase reporter plasmid or with pmirGLO the control plasmid (0.4 μg) using VIROMER^®^ RED reagent (Lipocalyx), according to the manufacturer’s instructions. Luciferase activity was determined after 48 h using a Dual Glo^®^ Luciferase Assay (Promega) and an Infinite M200 PRO luminometer (Tecan). Relative luciferase activities were calculated as the ratio of *Renilla* luciferase activity to firefly luciferase activity and further normalized to those of the control treatments.

## ACKNOWLEDGMENT

We would like to acknowledge Robert Pasieka for his input to the experimental part of this project and Susheel Bhat for his input during manuscript preparation. This work was supported by the Leading National Research Centre (KNOW) in Poznan under Grant 01/KNOW2/2014 (to PP, MS, AS, MZ, AP, and KDR). The research and related results of M.-D.R. were made possible by the support of the NOMIS Foundation. Deep sequencing was performed at the Laboratory of High Throughput Technologies (Faculty of Biology, Adam Mickiewicz University, Poznan) funded by National Multidisciplinary Laboratory of Functional Nanomaterials NanoFun nr POIG.02.02.00-00-025/09. The computations were partly performed at the Poznan Supercomputing and Networking Center.

