## Supplemental Information for "FUS controls the processing of snoRNAs into smaller RNA fragments that can regulate gene expression"

**MATERIALS AND METHODS**

The following primary antibodies were used in this work: anti-FUS (generated in-house (1)), anti-actin (MP Biomedicals, 691001), anti-Symplekin (Bethyl Laboratories, A301-465A), anti-FLAG (Sigma-Aldrich, A8592), anti-BRE (Santa Cruz Biotechnology, sc376453). The following secondary antibodies were used: goat anti-rabbit IgG-HRP and goat anti-mouse IgG-HRP (both from Santa Cruz Biotechnology).

**Cell culture and transfection**

HeLa, HEK293T and SH-SY5Y cells were grown in Dulbecco’s modified Eagle’s medium containing L-glutamine and 4.5 g*/*L glucose (DMEM; Lonza) and supplemented with 10% fetal calf serum (Gibco) and antibiotics (100 U*/*ml penicillin, 100 µg*/*ml streptomycin (Sigma-Aldrich)) at 37°C in a moist atmosphere containing 5% CO_2_. HEK293T cells with FUS OE, HEK293T cells with EBFP overexpression (EBFP OE) and HeLa cells with inducible FUS KD were prepared as described previously (1) (Supplementary Fig. S1). HeLa and SH-SY5Y cells with FUS KO were described previously (2, 3) (Supplementary Fig. S1). Plasmid DNA transfections (1 µg of plasmid) were performed with VIROMER® RED (Lipocalyx) and FuGENE® HD Transfection Reagent (Roche) according to the manufacturer’s instructions. For sdRNA104 5’ depletion, the improved method for ASO-mediated degradation of small nuclear RNAs in cultured mammalian cells, as described in (4), was used. The ASO was 20 nucleotides long, complementary to the full length of sdRNA104 5’, and consisted of 2’-O-methoxyribonucleotide segments of 5 nucleotides at each terminus and a deoxynucleotide segment containing 10 central nucleotides. The phosphate backbones were converted to phosphorothioate. ASOs were introduced at a concentration of 150 nM into 5 × 10^5^ HeLa cells using Lipofectamine 2000 reagent (Invitrogen, Thermo Fisher Scientific). Cells were collected 3 days later (Supplementary Fig. S3A).

**RNA isolation, cDNA preparation, PCR and qPCR**

RNA isolation, PCR and qPCR were performed as described in (1). The statistical significance of the qPCR results was determined by Student’s t-test. cDNA was synthesized using random hexamers or via a coupled polyadenylation reverse transcription reaction (to assess mature snoRNA and pre-snoRNA levels) (1). Stem-loop reverse transcription was performed with 200 ng of total RNA using a previously described protocol (5). To detect lariat structures, cDNA was amplified by PCR using outward-facing primers, and the detected product with the expected size was further cloned and sequenced, as described in (6). The primers used for PCR and qPCR are available on request.

**Protein isolation, RNA immunoprecipitation, and Western blotting**

Total protein extracts were prepared by lysis as described in (7) using a hypotonic lysis buffer or radioimmunoprecipitation assay (RIPA) buffer without SDS. For RNA immunoprecipitation, total and nuclear protein extracts from cells overexpressing FLAG-tagged FUS and EBFP proteins were gently rotated overnight at 4°C with anti-FLAG antibody-coupled magnetic beads (Sigma-Aldrich) and were then washed five times with PBS-T buffer. After washing, coprecipitated RNA was incubated with 1 U RNase T1 (Ambion) for 20 min at 25°C with shaking at 500 rpm. Next, samples were washed three times with PBS-T and treated with 50 U of alkaline phosphatase (Thermo Fisher Scientific) for 20 min at 25°C with shaking at 500 rpm to remove phosphate groups at the 3’ ends of RNA, followed by three more PBS-T washes. RNA was then treated with 20 U of kinase (OptiKinase, Affymetrix) to add phosphate groups at the 5’ ends. The samples were washed three times with PBS-T, and RNA was eluted from the beads with TRIzol reagent and used for library preparation. For Western blot analysis, samples were separated by sodium dodecyl sulfate-polyacrylamide gel electrophoresis (SDS-PAGE), transferred to polyvinylidene difluoride (PVDF) membranes (Millipore) and incubated first with specific primary antibodies and then with corresponding species-specific horseradish peroxidase (HRP)-conjugated secondary antibodies, followed by detection using enhanced chemiluminescence (ECL, GE Healthcare). The signals were quantified with Multi Gauge V2.2 software.

**Library preparation for RNA-seq**

Libraries were prepared using a TruSeq Small RNA Library Prep Kit (Illumina). Briefly, immunoprecipitated RNAs derived from particular samples were ligated to 3’ and 5’ RNA adapters. In further steps, reverse transcription and PCR were performed. PCR products were indexed via specific RNA PCR Index Primers (RPI, Illumina). PCR products were separated by electrophoresis on 6% polyacrylamide gels containing 1% glycerol. After 10 min of staining in SYBR™ Gold Nucleic Acid Gel Stain (Invitrogen, Thermo Fisher Scientific)/0.5x TBE buffer, DNA fragments of 140 bp-300 bp in length were cut and eluted overnight at 28°C with shaking at 400 rpm in 400 μl of elution buffer (50 mM Mg acetate, 0.5 M ammonium acetate, 1 mM EDTA, and 0.1% SDS). After chloroform/phenol purification, libraries were precipitated using 3 volumes of 100% EtOH in the presence of 1.5 μl of GlycoBlue™ coprecipitant (15 mg/ml) (Ambion, Thermo Fisher Scientific). Purified libraries were quantified using an Infinite® 200 PRO microplate plate reader with a NanoQuant Plate™ (Tecan) and Quant-iT™ PicoGreen™ dsDNA Assay Kit dye (Invitrogen, Thermo Fisher Scientific). Libraries were then pooled in equimolar ratios before sequencing. Sequencing was performed using a TruSeq SR Cluster kit v3-cBot-HS (Illumina), a TruSeq SBS kit v3-HS (50 cycles) (Illumina) and a HiScan™ SQ System platform (Illumina).

**Bioinformatic analysis**

The Illumina 3’ adapter sequences were trimmed from the raw sequencing reads using Cutadapt (available online at <http://journal.embnet.org/index.php/embnetjournal/article/view/200>). The quality of the cleaned libraries was verified using FastQC (available online at <http://www.bioinformatics.babraham.ac.uk/projects/fastqc>). Next, Bowtie (8) (adjusted to return only hits within the best strata) was used to align cleaned reads to the reference human genome assembly (GRCh37) obtained from the Ensembl database. SeqMonk software was used for quantification. Briefly, overlapping reads were joined into contigs, and the RPM (reads per million reads) value for every contig was calculated. Next, contigs were annotated with overlapping genomic features using the BEDTools suite and in-house scripts, allowing for multiple annotation of single contigs. Finally, contig enrichment in the immunoprecipitated sample was calculated as a log_2_-fold change over the control sample. Similarly, nuclear enrichment of the contigs was calculated as a log_2_-fold change over the total RNA sample.

Small RNA sequencing datasets were downloaded from Sequence Read Archive (9) (run IDs: SRR1586016 (HEK293, GSM1513689, 185.7 M bases); SRR3931966 (human neuroblastoma cells treated with sevoflurane, 661.6 M bases) and SRR2728234 (blood (PTSD with comorbid depression, GSM1912509, 683 M bases). The reads were subjected to adapter clipping with FASTX-Toolkit (<http://hannonlab.cshl.edu/fastx_toolkit/>), and only reads corresponding to adapter-containing small RNAs > 18 bases long were kept. Then, reads were mapped to a set of human snoRNA sequences from Ensembl (10) with Bowtie. The mappings were then subjected to manual inspection to keep only putative snoRNA-derived miRNAs (length ≤ 24 nt, raw count > 10, most abundant per sequence arm) and the most abundant sequences longer than 24 bases and a raw count of > 50, representing the most abundant mapped reads per sequence arm. To predict and analyze targets for short and long (>24 nt) sRNAs, Miranda (11) was used with two approaches: i) using the whole transcriptome and ii) using only the 3' UTRs of the human transcripts. The human transcriptome was downloaded from Ensembl 86 (10). Then, the top 100 genes with the highest Miranda scores (i.e., those predicted to form the highest number of and/or most stable interactions with sRNAs) were selected. The gene IDs obtained via this approach were then used as the input to overrepresentation analysis for pathways and gene ontologies using ConsensusPathDB (<http://consensuspathdb.org/>) (12). The predicted sRNA target regions were then compared to those in HEK293T-specific data from photoactivatable ribonucleoside-enhanced crosslinking and immunoprecipitation (PAR-CLIP) and high-throughput sequencing of RNA isolated by crosslinking immunoprecipitation (HITS-CLIP) experiments for AGO1, AGO2, AGO3 and AGO4 from the StarBase 2.0 database (13) after converting the genome coordinate system from hg19 to hg38 using LiftOver (<https://genome.ucsc.edu/cgi-bin/hgLiftOver>). This step was done with the Intersect tool from the BEDTools suite (14), requiring that at least 50% of the targeted region fell in the CLIP-Seq cluster. Only Miranda sRNA target predictions that overlapped CLIP-Seq peaks were kept for further consideration.

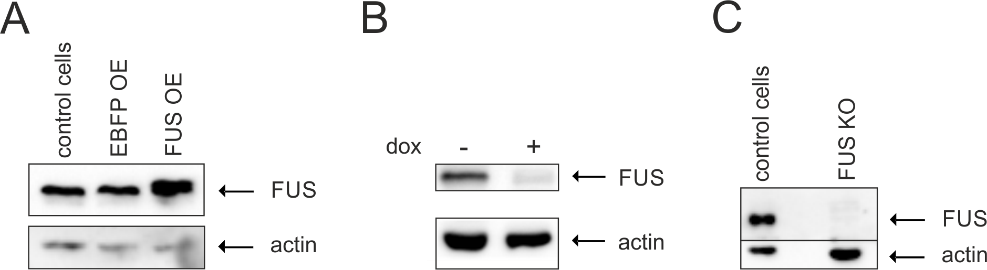

*Supplementary Figure S1. Western blotting followed by immunodetection with anti-FUS and anti-actin antibodies was performed using protein extracts from A) wild-type cells (control cells), cells with EBFP or FUS overexpression (EBFP OE or FUS OE, respectively), B) cells with inducible FUS knockdown with (dox+) or without (dox-, control cells) doxycycline treatment and C) cells with FUS knockout (FUS KO).*

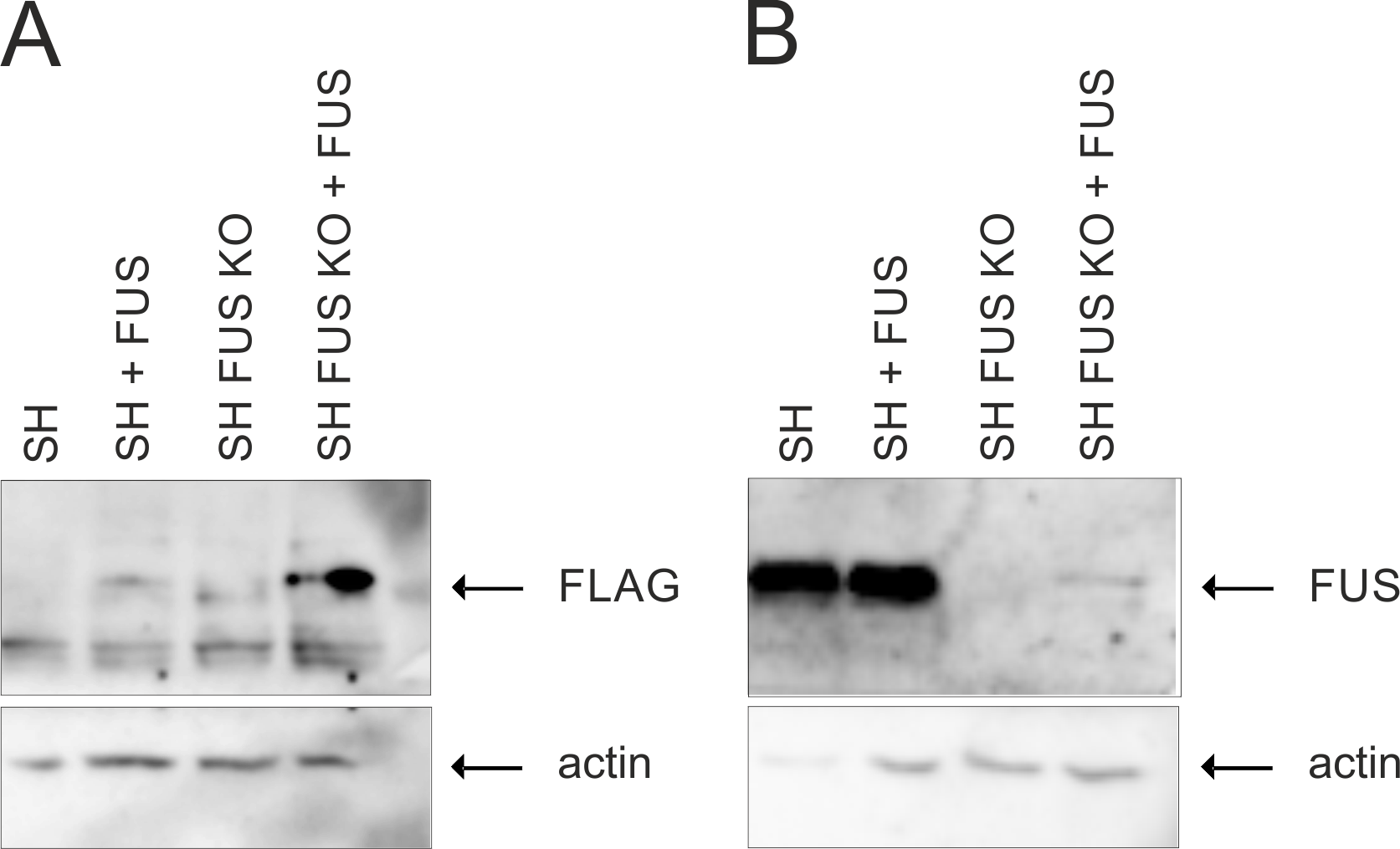

*Supplementary Figure S2. Western blotting followed by immunodetection with A) anti-FLAG and B) anti-FUS and anti-actin antibodies was performed using protein extracts from wild-type SH-SY5Y cells (SH), SH-SY5Y cells with transient overexpression of FUS (SH + FUS), SH-SY5Y cells with FUS knockout (SH FUS KO) and SH FUS KO cells with transient overexpression of FUS (SH FUS KO + FUS).*

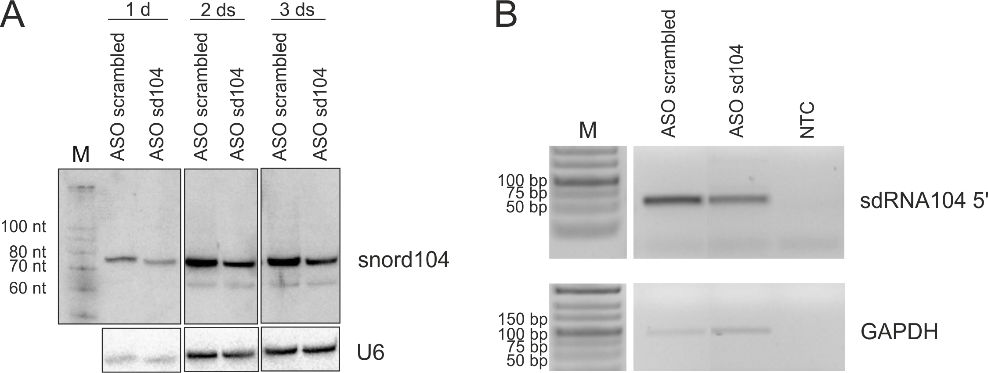

*Supplementary Figure S3. Northern blotting and SL RT-PCR were performed to confirm decreased levels of sdRNA104 in cells in which sdRNA104 was knocked down using an ASO strategy. A) Northern blots with probes targeting sdRNA104 5’ and U6 snRNA were performed using RNA extracts from cells transfected with scrambled ASO or sdRNA104 ASOs for 1 day, 2 days and 3 days. B) SL RT-PCR was performed using cDNA prepared from cells transfected with scrambled ASO or sdRNA104 ASO for 3 days; NTC–nontemplate control.*
